# Praziquantel activates a native cation current in *Schistosoma mansoni*

**DOI:** 10.1101/2023.08.25.554840

**Authors:** Evgeny G. Chulkov, Claudia M. Rohr, Jonathan S. Marchant

**Affiliations:** Department of Cell Biology, Neurobiology, and Anatomy, Medical College of Wisconsin, Milwaukee, WI, USA

**Keywords:** parasite, anthelmintic, schistosome, ion channel, TRP channel

## Abstract

Praziquantel (PZQ), an anthelmintic drug discovered in the 1970s, is still used to treat schistosomiasis and various other infections caused by parasitic flatworms. PZQ causes a triad of phenotypic effects on schistosome worms – rapid depolarization, muscle contraction, and damage throughout the worm tegument. The molecular target mediating these effects has been intimated as a Ca^2+^-permeable ion channel, but native currents evoked by PZQ have not been reported in any schistosome cell type. The properties of the endogenous PZQ activated conductance therefore remain unknown. Here, invasive electrophysiology was used to probe for responses to PZQ from different locales in a living schistosome worm. No direct response was seen in tegument-derived vesicles, or from the sub-tegumental muscle layer despite the presence of voltage-operated currents. However, PZQ rapidly triggered a sustained, non-selective cation current in recordings from neuronal tissue, targeting both the anterior ganglion and the main longitudinal nerve cord. The biophysical signature of this PZQ-evoked current resolved at single channel resolution matched that of a transient receptor potential ion channel named TRPM_PZQ_, recently proposed as the molecular target of PZQ. The endogenous PZQ-evoked current was also inhibited by a validated TRPM_PZQ_ antagonist. PZQ therefore is a neuroactive anthelmintic, effecting a robust, depolarization through ion channels with the characteristics of TRPM_PZQ_.

**Key Findings / Scope Statement:**

- Responses to the anthelmintic drug, praziquantel (PZQ), were examined using invasive electrophysiology in a living schistosome worm.
- PZQ evoked a cation current in recordings from neuronal tissue
- The biophysical and pharmacological characteristics of the native PZQ current matched the properties of TRPM_PZQ_.

## Introduction

The study of excitable cell physiology in parasitic flatworms has long been a focus for research (Geary et al., 1992, Pax et al., 1996, Greenberg, 2014, McVeigh et al., 2018). This is because of the likelihood for discovering vulnerabilities to chemotherapeutic attack within the transmembrane signaling portfolio of these cells. Many existing anthelmintic agents are known to subvert targets that control parasite neuronal and/or muscular function.

One such example is the drug praziquantel (PZQ), the key clinical drug used to combat schistosomiasis. PZQ causes a spastic paralysis of schistosome musculature by stimulating rapid depolarization and Ca^2+^ entry that effects a sustained, tetanic increase in muscle tension (Andrews et al., 1983, Park and Marchant, 2020, Waechtler et al., 2023). This activity is widely seen in different parasitic flatworms that are sensitive to PZQ, and is blocked by removal of Ca^2+^, or application of certain Ca^2+^ channel blockers (Pax et al., 1978, Fetterer et al., 1980a). These observations have long supported a ‘Ca^2+^ channel activation’ hypothesis for PZQ action (Jeziorski and Greenberg, 2006, Chan et al., 2013). However, the molecular basis for these effects has long proved elusive, with no endogenous target for PZQ unmasked throughout decades of clinical usage. Such lack of insight has been exacerbated by an inability to resolve any native current evoked by PZQ in schistosomes, or indeed any parasitic flatworm.

The majority of our knowledge about endogenous ion channel function in schistosomes derives from pioneering experiments performed in the 1980s and 1990s which resolved fundamental features of voltage gradients in native worms (Fetterer et al., 1980b, Bricker et al., 1982, Semeyn et al., 1982, Thompson et al., 1982), with examples of electrophysiological recordings from isolated muscle cells (Blair et al., 1991, Day et al., 1993, Day et al., 1995, Mendonca-Silva et al., 2006), tegument (Day et al., 1992) and tegument-derived vesicles (Robertson et al., 1997). These assays lead to the description of several different types of ion fluxes, including currents mediated by Cl^-^ channels (Robertson et al., 1997), voltage-operated Ca^2+^ channels (Mendonca-Silva et al., 2006), various K^+^ channels (Day et al., 1993, Day et al., 1995, Kim et al., 1995a, Kim et al., 1995b, Robertson et al., 1997), a Ca^2+^-activated K^+^ channel (Blair et al., 1991), and other non-selective cation channels (Day et al., 1992, Robertson et al., 1997).

Despite such efforts, a native response to PZQ remained either unresolved or unreported. A possible reason, beyond the technical challenge of measuring native currents from parasitic flatworms, was the lack of insight as to what exactly to look for, and where exactly to look. Additionally, in the absence of any knowledge about the characteristics of the target, the specific recording conditions to best resolve PZQ-evoked endogenous currents remained undefined.

Recent advances have however increased the temptation to have another stab at this challenge. First, a candidate target for PZQ has been identified – an ion channel of the transient receptor potential melastatin family, named TRPM_PZQ_ (Park et al., 2019, Park and Marchant, 2020). Second, identification of this target provides direction as to where to look for native currents - based on the atlas of single cell RNA expression data in schistosomes, *Schistosoma mansoni* TRPM_PZQ_ (*Sm*.TRPM_PZQ_) is expressed in several neuronal and muscle clusters (Wendt et al., 2020). Third, electrophysiological analyses of *Sm*.TRPM_PZQ_ have now been executed (Park et al., 2021, Chulkov et al., 2023b), establishing a search algorithm for the likely PZQ-evoked response, as well as conditions best optimized to resolve TRPM_PZQ_ currents. *Sm*.TRPM_PZQ_ is a non-selective cation channel with a linear current-voltage relationship (Chulkov et al., 2023b)). Of relevance here, *Sm*.TRPM_PZQ_ display a clear permeability toward Cs^+^, and this provides opportunity to record currents in the absence of many types of K^+^ channels. Finally, recent drug screening efforts have yielded antagonists that block *Sm*.TRPM_PZQ_ activity (Chulkov et al., 2021). Capitalizing upon all this new information, electrophysiological recordings were attempted from different types of tissue within a living worm. An endogenous PZQ-activated current was identified in recordings from putative neuronal locales, the biophysical characteristics of which resembled the properties of *Sm*.TRPM_PZQ_.

## Materials and Methods

### Materials

All chemicals were sourced from Sigma or ThermoFisher. Praziquantel was used as a racemic mixture ((±)-PZQ). ANT1 was sourced from Maybridge (Chulkov et al., 2021).

### Adult schistosome worm isolation

Schistosome-infected mice (*Schistosoma mansoni*) were provided by the NIAID Schistosomiasis Resource Center at the Biomedical Research Institute (Rockville, MD) through NIH-NIAID Contract HHSN272201000005I for distribution via BEI Resources. Adult schistosomes were recovered by dissection of the mesenteric vasculature in female Swiss Webster mice previously infected (∼49 days) with *S. mansoni* cercariae (NMRI strain). All experiments followed ethical regulations endorsed by the Medical College of Wisconsin IACUC committee. Harvested worms were washed in DMEM high glucose medium, supplemented with HEPES (25mM), pyruvate and 5% heat inactivated FBS (Gibco) and penicillin-streptomycin (100 units/mL) and incubated overnight (37°C/5% CO_2_) in vented petri dishes (100×25mm).

### Electrophysiological assays

Electrophysiological assays were then performed over a period of four days following worm isolation. For these assays, a single, male, adult schistosome was either pinned with dual needles, or fixed with glue (n-butyryl 3M Vetbond™ surgical glue; 3M, St. Paul MN) onto a 90mm Sylgard™-coated plastic dish (Living Systems, St Albans VT). For recordings from tegument (Figure 1), muscle (Figure 2) and lateral nerve cords (Figure 4), the adult worm was pinned to avoid any pervasive damage across the worm surface. For recordings from anterior neurons (Figure 3), where a high degree of immobility was required for successful recordings, the dorsal surface was glued to the dish. Access to the anterior region of the worm was facilitated by immobilizing worms in this manner. Recordings were only made from male worms, owing to their greater size that better facilitated invasive electrophysiology.

**Figure 1.**
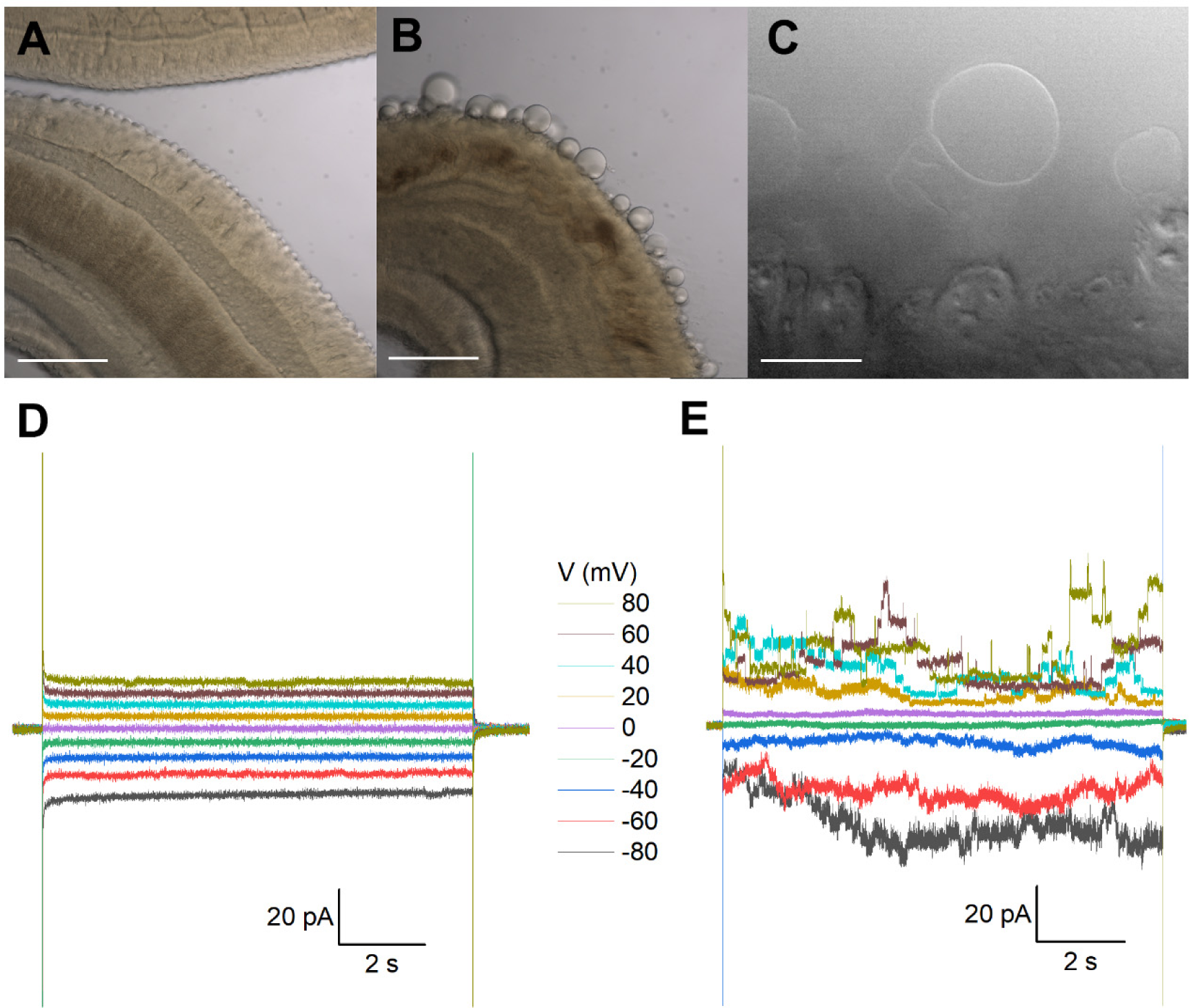
Recordings from tegumental vesicles derived by treatment of *S. mansoni* with PZQ. (**A**) Brightfield image showing a region of an adult, male *S. mansoni* worm under control conditions. (**B**) Brightfield image of an adult, male *S. mansoni* worm after treatment with PZQ (5 µM, 24 hr) to illustrate blebbing of the tegument. Scalebar, 200 µm (A&B). (**C**) Examples of giant unilamellar vesicles formed from the worm tegument after exposure of an adult *S. mansoni* worm to PZQ (10 µM, 15 min). Scalebar, 50 µm. (**D**) Representative current traces from a vesicle-attached patch at different voltages. (**E**) Representative current traces from a cell-attached patch from a HEK293 cell co-transfected with *Sm*.TRPM_PZQ_ and GFP at different voltages. In all recordings in this figure, bath solution: HBSS with PZQ (10 µM); pipette solution, 140mM CsMeSO_3_, 10mM HEPES, 1mM EGTA, pH 7.4.

**Figure 2.**
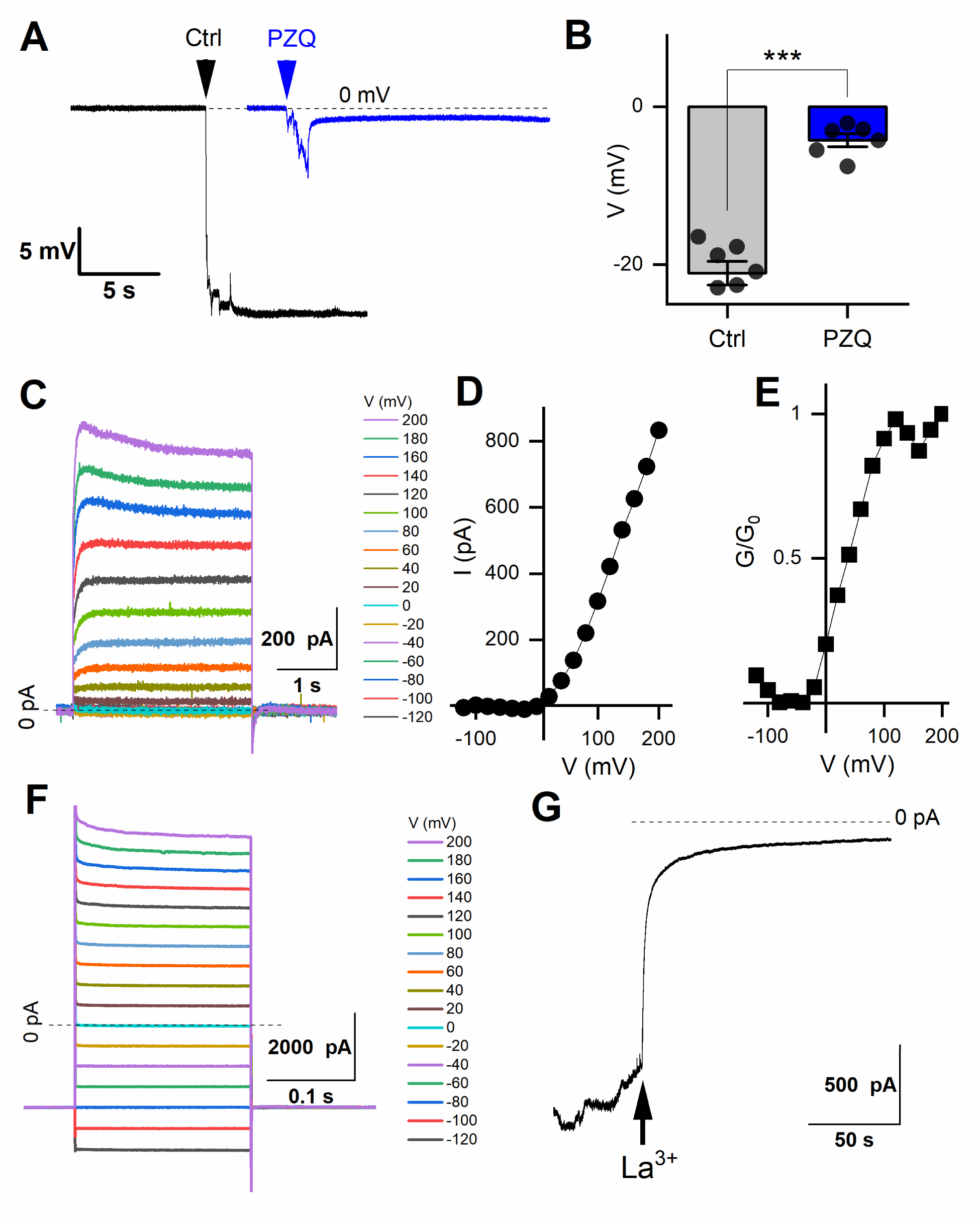
Recordings from schistosome muscle. (**A**) Representative traces of electrical potentials (mV) registered upon penetration of the dorsal surface of an adult schistosome worm under either control (‘Ctrl’) conditions (vehicle, 0.1% DMSO) or in the presence of PZQ (10 µM in bath solution, ‘PZQ’). Arrow indicates the moment of electrode penetration. With the electrode positioned in the bath a stable potential (0 mV) was resolved before penetration into the muscle layer. Bath solution: HBSS (20 mM HEPES, 100 µM carbachol, pH 7.4). Pipette solution: (3M KCl). (**B**) Steady-state value of the peak electrical potential recorded (mean±SE, n=6 worms for each group) in these assays (***, p<0.001). (**C**) Representative current traces recorded from a worm muscle at different voltage steps from a holding voltage of −80mV. (**E**) Normalized slope conductance (G/G_0_, where G is conductance at a specific voltage and G_o_ is the maximum slope conductance) versus voltage plot from worm muscle recordings. (**F**) Representative current traces from worm muscle at different voltage steps from a holding voltage of −80mV recorded with PZQ in the bath solution. (**G**) Representative current trace showing blockade of currents in worm muscle exposed to PZQ following addition of 5 mM LaCl_3_ (arrow) to the bath. Holding voltage −80 mV. For all experiments in this figure, recordings were made in bath solution: HBSS with 100μM carbachol; pipette solution: 140mM CsMeSO_3_, 10mM HEPES, 1mM EGTA, pH 7.4.

**Figure 3.**
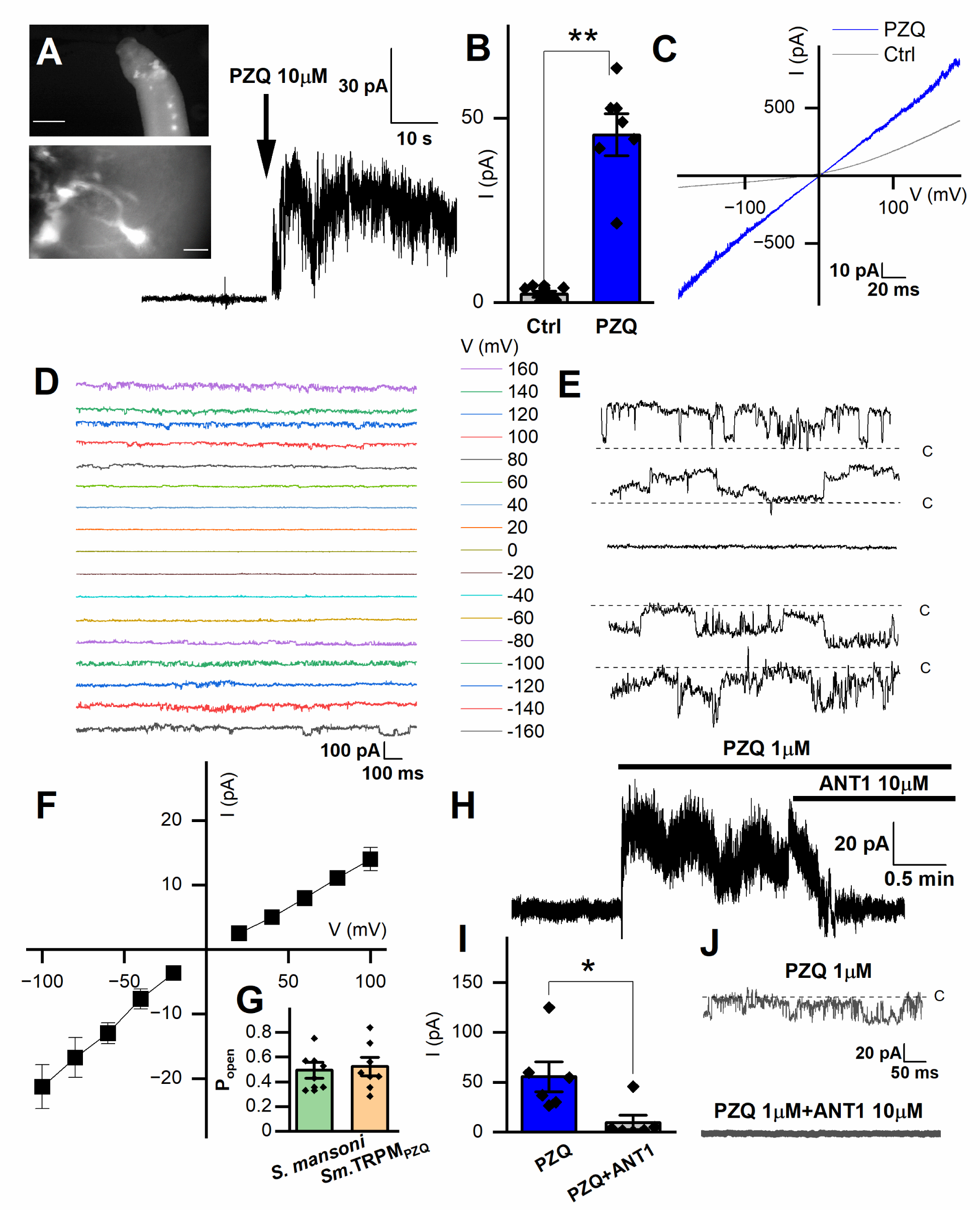
Records targeting the anterior ganglion. (**A**) *Inset top*, image showing fluo-4 fluorescence resolved in the anterior region of a male, adult schistosome worm. Scalebar, 500µm. *Inset bottom*, enlarged image of an example cell body with extended processes. Scalebar, 10µm. Representative recording from an adult male *S. mansoni* anterior neuron prior to and after addition of PZQ (10µM, *arrow*). (**B**) Maximum plateau current (I, pA) before (‘Ctrl’) and after addition of PZQ (10µM), ** p≤0.01, n≥6. (**C**) Representative current (I) – voltage (V) plot in the absence (‘Ctrl’) and in the presence of PZQ (10 µM). (**D**) Representative current traces recorded at different voltages from an anterior neuron patch in the presence of PZQ (10µM). (**E**) Single channel like fluctuations recorded from an anterior worm neuron in the presence of PZQ (10µM) at 120 mV (top), 80 mV, 0 mV, −80 mV and −120 mV (bottom). (**F**) Current (*I*) – voltage (*V*) plot from single channel like unitary currents recorded from the anterior worm ganglion. (**G**) Open probability (P_open_) of single channel fluctuations compared between *S. mansoni* and HEK293 cells expressing *Sm.*TRPM_PZQ_ in the presence of PZQ (10µM) in the bath solution. (**H**) Representative current trace from an anterior *S. mansoni* neuron before and after addition of PZQ (1µM), and then addition of ANT1 (10µM). (**I**) Mean steady current in the presence of PZQ (1µM) or in the presence of PZQ (1µM) and ANT1 (10µM). p≤0.05, n≥6. (**J**) Representative current trace from worm neurons at −80 mV in the presence of PZQ (1µM) or PZQ (1µM) and ANT1 (10µM). For all experiments in this figure, recordings were made in bath solution: HBSS; pipette solution: 140mM CsMeSO_3_, 10mM HEPES, 1mM EGTA, pH 7.4.

**Figure 4.**
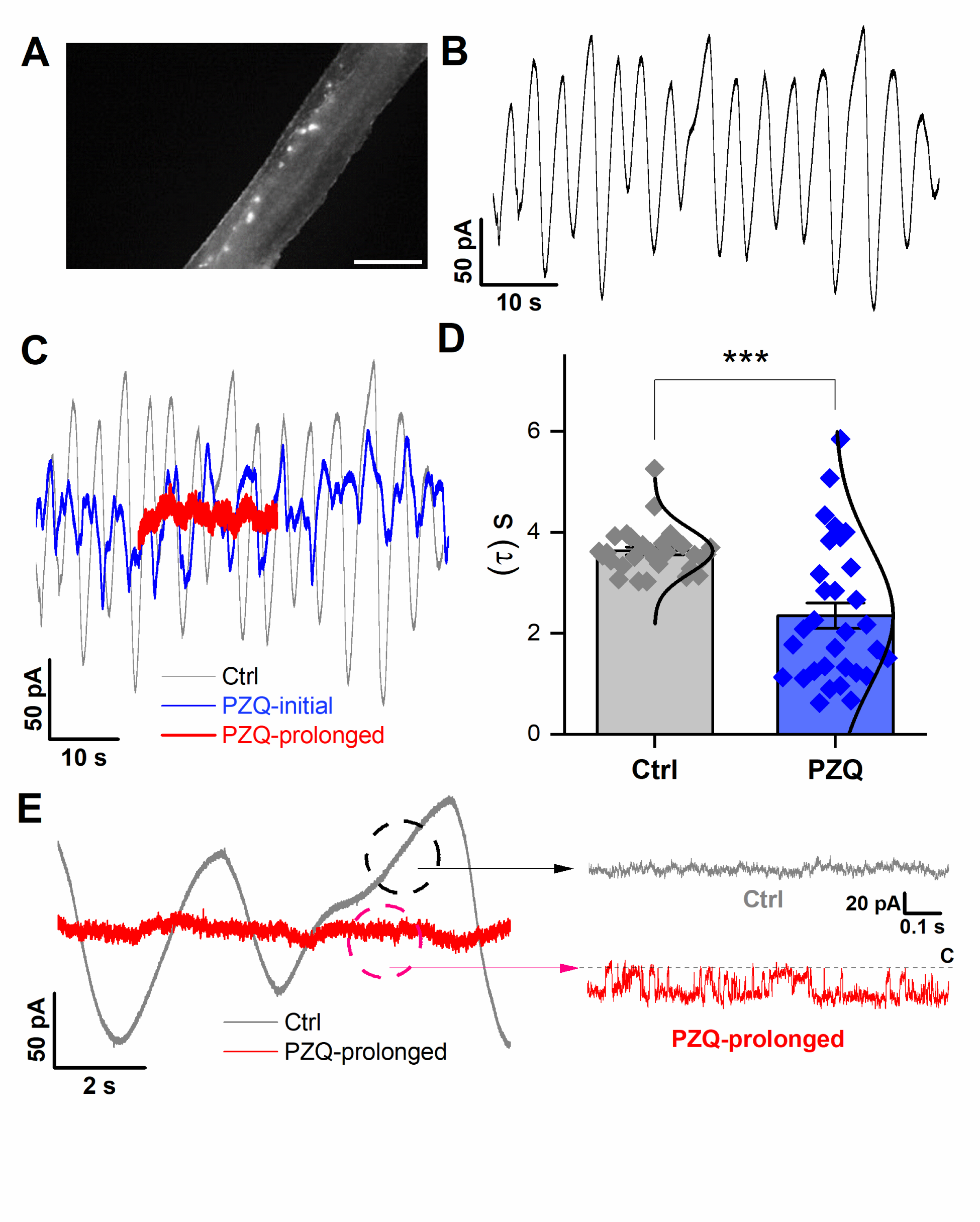
Recordings targeting the longitudinal nerve cords. (**A**) Image showing fluo-4 fluorescence resolved along the body of a male, adult schistosome worm. Scalebar, 500µm. (**B**) Representative current oscillation recorded from the lateral nerve cord resolved after a successful seal is made into neuronal tissue. (**C**) Effect of PZQ (10µM) on the endogenous oscillation evident by comparing the waveform before (‘Ctrl’), immediately after (‘PZQ-initial’, blue) and after prolonged (>5min, red) drug exposure (‘PZQ-prolonged’). (**D**) Peak-to-peak period (τ, sec) of the current oscillations recorded in the absence (‘Ctrl’) or after addition of PZQ (10µM). Data are from three separate recordings in different worms. (**E**) Representative trace of current from worm lateral cord neurons recorded at −160 mV either in the absence (‘Ctrl’, slope baseline subtracted) or prolonged presence of 10µM PZQ (‘PZQ-prolonged’). For all experiments in this figure, recordings were made in bath solution: HBSS; pipette solution: 140mM CsMeSO_3_, 10mM HEPES, 1mM EGTA, pH 7.4.

Recording dishes were mounted on the stage of an Olympus BX51WI upright microscope. Electrodes were pulled from borosilicate glass capillaries (#BF150-110-10, Sutter Instrument, Novato, CA) on a vertical pipette puller (Narishige, Amityville, NY, Model PC-10). Except where specified otherwise, the bath solution was Hank’s balanced salt solution (HBSS) supplemented with 20 mM HEPES (pH 7.4 at room temperature). For recordings from neuronal tissue, worms were loaded with a Ca^2+^ indicator (fluo-4-NW dissolved in HBSS with 2.5 mM probenecid) by incubation for 30 mins at 37°C, and fluorescence was visualized by epifluorescence illumination using a Spark camera (Hamamatsu, Japan).

A multiClamp 700B amplifier and Digidata 1440A digitizer (Molecular Devices, San Jose, CA) were used for electrophysiological recordings. Signals were passed through an 8-pole Bessel low pass filter at 1 kHz and sampled at 10 kHz. Data analysis was performed using Clampfit 11 software (Molecular Devices). In current-clamp recordings, microelectrodes (resistance of 2 – 4 MΩ) filled with 3M KCl were advanced using a micromanipulator, penetrating the worm surface, until a steady signal was obtained in the current-clamp mode (Thompson et al., 1982). For muscle voltage measurements, the bath solution was supplemented with 100 µM carbachol to impair muscle contraction (Barker et al., 1966, Thompson et al., 1982). Electrical potentials were measured after penetrating the dorsal tegument at an angle of 30° from horizontal. Distinctive steady-state voltages were observed depending on the depth of recording within internal schistosome tissue (Bricker et al., 1982, Thompson et al., 1982). In voltage-clamp mode, current measurements were performed with electrodes pulled to a resistance of 8 – 10 MΩ). The pipette solution used throughout was 140mM CsMeSO_3_, 10mM HEPES and 1mM EGTA (pH 7.4). Recordings from HEK293 cells were performed as detailed previously (Chulkov et al., 2023a).

### Statistical Analyses

Data were visualized and processed using Origin (2020b). Tukey’s test was applied to evaluate significance between different cohorts of measurements, and reported as mean and standard error of the mean (mean ± SE). The number of recordings associated with individual experiments are defined in the Figure legends.

## Results

Attempts to measure native currents in schistosomes were made by electrophysiological recording from different locales within a living worm. First, recordings were made from tegument-derived vesicles. Second, an invasive electrophysiology approach was used to resolve responses from the muscle layer localized beneath the tegument. Finally, invasive recordings were attempted from nervous tissue. These encompassed measurements targeting either the anterior cephalic ganglion, or the main longitudinal nerve cords. Details of each of these approaches are outlined in the following sections.

### Recordings from tegument-derived vesicles

The surface of schistosomes comprises a living syncytium known as the tegument that is bounded by a double outer bilayer, insulating the parasite from the host bloodstream (Wilson and Jones, 2021). While this surface is readily accessible, electrophysiological recording is difficult owing to the rough and spiny nature of the male tegument. This precluded formation of a tight seal (giga-ohm resistance) necessary for single channel recording. However successful recordings have previously been achieved by clamping vesicles derived from the tegument. One method for generating vesicles is exposure of worms to a low pH (pH ∼3.75), a manipulation that yielded a population of smooth, bilayered structures (Robertson et al., 1997). Treatment with PZQ also causes extensive vesicularization at the worm surface (Becker et al., 1980), providing an alternative method to generate ‘worm-free’, self-formed vesicles accessible to electrophysiological analysis.

Treatment of male worms with PZQ (10 µM, 15 mins) caused surface vesicularization (**Figure 1A & B**), generating unilamellar lipid vesicles that were easily visualized using bright-field illumination (**Figure 1C**). It was facile to form a tight seal (>1GΩ) onto these vesicles with the recording electrode permitting ‘vesicle-attached’ recordings. However, using this preparation, no responses to PZQ application were observed throughout a range of voltage-steps or recording conditions (**Figure 1D**). Given the tight seal between the recording electrode and membrane, a channel such as *Sm*.TRPM_PZQ_ (conductance of ∼130pS in symmetrical 145mM Na^+^, (Chulkov et al., 2023b)) would be associated with fluctuations of up to 10 pA. For those unfamiliar with electrophysiological approaches, a similar voltage-stepping protocol in *Sm*.TRPM_PZQ_ expressing HEK293 cells is illustrated for comparison (**Figure 1E**). Here, clearly resolvable ion channel activity was seen following addition of PZQ.

### Recordings from the internal muscle layer

Layers of circular, longitudinal, and transverse smooth muscle fibers exist below the adult schistosome tegument (Silk and Spence, 1969, Sulbaran et al., 2015) The electrical properties of this tissue layer have been investigated by invasive recording approaches, where discrete potential changes are observed when the recording electrode penetrates different compartments of the worm (Bricker et al., 1982, Thompson et al., 1982). For example, in current clamp mode (I=0pA), a drop in potential of ∼25-30mV is associated with penetration into sub-tegumental muscle. A similar drop in voltage (−21.1 ± 3.9mV) was seen in our recordings when the recording electrode penetrated into the tissue immediately beneath the tegument (**Figure 2A**). In worms treated with PZQ (10 µM, 10 minutes), this drop in membrane potential was no longer resolved, with only a small voltage change (−4.2 ± 2.0 mV) observed on electrode penetration (Figure 2A). **Figure 2B** collates the electrical potentials observed in either control (−21.1 ± 3.9mV) or PZQ-treated (−4.2 ± 2.0 mV) worms. The existence of the potential in naïve worms, and the loss of this gradient after incubation with PZQ, is characteristic of recordings being made from sub-tegumental muscle (Bricker et al., 1982). That recordings at this depth beneath the tegument were from muscle tissue was further evidenced by measurements made in voltage-clamp mode. At this depth within the worm body, voltage-dependent currents were observed (**Figure 2C**). With the membrane held at −80 mV, various voltage pulses (−120mV to 200mV) were applied. Currents were seen at positive, but not at negative membrane voltages in naïve worms (**Figure 2D**), reaching a peak conductance ratio (G/G_0_) at ∼100 mV of applied voltage (**Figure 2E**). PZQ treatment (10 µM, 10 minutes) was associated with an increase in overall ionic permeability in muscle, and voltage-dependent responses did not persist after PZQ treatment (**Figure 2F**). The existence of increased cation currents in muscle in the presence of PZQ was shown by addition of La^3+^ (5mM), which blocked these currents (**Figure 2G**). These data show that PZQ impacts schistosome muscle physiology, causing maintained depolarization associated with a loss of voltage-dependent ion channel activity.

### Recordings from neural tissue

In terms of neuroanatomy, the central nervous system of adult schistosomes consists of a bi-lobed, anterior cephalic ganglion from which pairs of longitudinal nerve cords project and run the length of the worm (Hyman, 1951, Halton and Gustafsson, 1996, Halton and Maule, 2004). These longitudinal nerve cords are cross-linked by transverse fibers along the body axis. A peripheral nervous system also interconnects the major body organs. Visualization of these structures in a living worm is challenging in the absence of a selective labelling method. However, we noted that incubation (∼30 mins) of intact worms with the Ca^2+^-sensitive dye fluo-4 (fluo-4-NW) resulted in compartmentalization of the fluorescent indicator within structures apparent both anteriorly and in tracts running longitudinally down both sides of the worm (Figure 3 & Figure 4). The concentration of dye, above levels of fluorescence staining apparent in surrounding tissues, was suggestive of neuronal structures with the region of anterior fluorescence likely representing the cephalic ganglion and the lateral structures of elevated fluorescence intensity reminiscent of the longitudinal nerve cords.

**Figure 3A** (top panel) shows examples of the anterior fluo-4 staining, with several cell bodies visible in the enlarged panel (Figure 3A, bottom). Using the axial distribution of fluo-4 fluorescence as a guide, the recording electrode was positioned with a micromanipulator into close proximity to where fluorescence was resolved. Voltage steps of 10mV (at a holding voltage of −80mV) were repeated at a frequency of 50Hz while the recording electrode was incrementally advanced. When a change in resistance occurred, negative pressure (10-20 mmHg) was applied, with a loose seal (0.2-0.4 GΩ) being formed in a minority of attempts (equating to a 10-20% success rate). Figure 3A shows a representative current trace recorded from one putative anterior neuron, where addition of PZQ (10µM, at 40 mV) activated a sustained inward current. Figure 3B compares the peak current before (2.3 ± 0.8pA) and after PZQ (46 ± 6pA).

Application of a voltage ramp (10 mV/s) after successful seal formation revealed a slight voltage dependence in the clamped cell in the absence of PZQ (Figure 3C). After addition of PZQ, currents were larger and linear, as the voltage-dependence was lost. Pronounced noise was evident at higher voltages (Figure 3C), suggesting the presence of single channel currents in the membrane. To investigate this further, we applied prolonged voltage steps in the presence of PZQ (10µM, holding voltage of 0 mV, Cs^+^ as the permeant inward cation, Figure 3D). The resulting current traces revealed single channel-like events at different voltages (Figure 3E). At 0 mV, no channel like fluctuations were observed (enlarged in Figure 3E). However, at larger positive and negative applied voltages, recognizable single channel like-activity was evident. The single channel current-voltage (I-V) plot was fitted with a linear regression, giving an estimated conductance of 154 ± 7 pS (Figure 3F). The open state probability of the endogenous PZQ-evoked channel fluctuations (P_open_ = 0.49 ± 0.06) (Figure 3G, recorded at 80 mV) were not statistically different from recordings made in *Sm*.TRPM_PZQ_ expressing HEK293 cells (P_open_ = 0.52 ± 0.07).

A recent target-based screen identified an antagonist (ANT1, 1-(9*H*-fluoren-9-yl)-4-(5-methyl-3-phenyl-1,2-oxazole-4-carbonyl)piperazine) of *Sm*.TRPM_PZQ_ that blocked PZQ-evoked channel activation and worm contraction (Chulkov et al., 2021). Addition of ANT1 (10µM) to the bath solution decreased the PZQ-activated neuronal current (Figure 3H). Cumulative measurements of peak current prior to, and after, ANT1 addition are shown in Figure 3I. The endogenous single channel-like fluctuations evoked by PZQ in worm neurons were also inhibited by addition of ANT1 (10µM) (Figure 3J). The biophysical and pharmacological signature of the *in vivo* response therefore resembles the properties of *Sm*.TRPM_PZQ_ measured *in vitro*

Finally, recordings were attempted from the lateral nervous plexus visualized targeting the fluo-4 fluorescence apparent along the longitudinal axis of the worm (Figure 4A). These recordings were especially challenging given the smaller area of fluorescence (success rate <5%). Following a successful seal onto ‘excitable’ tissue, an endogenous, slow oscillatory behavior was observed (Figure 4B). In naïve conditions, the current oscillations rapidly and regularly transitioned from minimum to maximum amplitude with a regular period of 3.6±0.1s (Figure 4B). Application of PZQ (10 µM) caused progressively increasing noise, suppressing the amplitude, and disrupting the period of the oscillations resulting in greater irregularity (Figure 4C). This was reflected in a broadening of the wave period distribution (Figure 4D). Prolonged exposure to PZQ (∼ 5 mins) caused the oscillations to cease (Figure 4C, red).

In the native state, during the endogenous current oscillations, scrutiny of the linear rising phase of the oscillation revealed no recognizable single channel activity (Figure 4E). However, evaluation of the current traces after prolonged PZQ exposure revealed distinguishable step-like fluctuations (Figure 4E). These signals, recorded at −160mV, exhibited a single channel current of 19.1±5.4pA and an open probability, P_open_ = 0.53 ± 0.17. These values are consistent with the properties displayed by *Sm*.TRPM_PZQ_ measured in HEK293 under similar recording conditions (Chulkov et al., 2023a, Chulkov et al., 2023b). Therefore, recordings from two different neuronal locales evidenced clear single channel activity in response to the application of PZQ. The properties of this native response were consistent with *Sm*.TRPM_PZQ_.

## Discussion

Here, we were able to resolve a native PZQ-evoked current in recordings from a live schistosome worm. To the best of our knowledge, this represents the first report of an endogenous ion channel current activated by PZQ in any parasitic flatworm. This current was devolved at single channel resolution, and the biophysical properties of this response (Cs^+^ permeability, P_open_ and the linear I-V relationship matched the *in vitro* electrophysiological characteristics of the ion channel, *Sm*.TRPM_PZQ_, which has been proposed as the parasite target of this drug (Park et al., 2019). However, the estimated conductance of the native PZQ channel in anterior neurons was 154 ± 7 pS (Figure 3F). This value is higher than the measured conductance of *Sm*.TRPM_PZQ_ (112 ± 12 pS) in a similar solution (140mM Cs+) after heterologously expression in HEK293 cells (Chulkov et al., 2023a). This difference may be due to the different lipid/intracellular ion composition of schistosome neurons versus human HEK293 cells, and the presence of additional outward currents in the worm neuronal background. The native response to PZQ was blocked by ANT1 (Figure 3), a validated antagonist of *Sm*.TRPM_PZQ_ (Chulkov et al., 2021). Overall, the biophysical and pharmacological properties of the ion channel underlying the native response to PZQ are comparable with known properties of *Sm*.TRPM_PZQ_.

PZQ-evoked currents were resolvable from two types of neuronal tissue (recordings that targeted either the anterior ganglion or lateral nerve cords). This is consistent with single cell RNAseq data localizing *Sm*.TRPM_PZQ_ expression to various types of neurons (Wendt et al., 2020). Therefore, these data are consistent with a model where PZQ acts directly on TRPM_PZQ_ expressed in neurons to effect neuronal depolarization. The lack of desensitization of TRPM_PZQ_ in response to PZQ would ensure a long-lasting neuronal depolarization, a sustained release of neurotransmitters and thereby protracted paralysis of muscle tissue. A sustained PZQ-evoked depolarization of muscle is consistent with the loss of voltage-sensitivity of currents in muscle after PZQ treatment (Figure 2). This excitotoxic tsunami could also underpin damage to the tegument, analogous to an inflammatory response in skin that is triggered by stimulus-evoked neurotransmitter release from sensory neurons. TRPM channels are well known to regulate exocytosis in various mammalian cell types (Brixel et al., 2010, Held et al., 2015).

This is the first application of invasive electrophysiology in parasitic flatworms. However, dissected preparations have previously been optimized to allow resolution of single-channel responses in various parasitic nematodes (Qian et al., 2006, Robertson et al., 2011). Here, the challenges inherent to the invasive electrophysiology approach merit a few caveats.

First, sampling bias. Despite efforts to record at different locations in the worm and at various depths of recording electrode penetration, this method is of course not a comprehensive analysis of all types of cells present in the worm. For example, while we did not observe a *direct* effect of PZQ on worm muscle cells, it is feasible that these recordings have not captured the needed diversity of different muscle cell types. If TRPM_PZQ_ is expressed only in a subtype of muscle cells (as suggested by single cell RNAseq studies (Wendt et al., 2020)), such cells may not have been sampled by our assays. However, noting this qualification, TRPM_PZQ_ single channel activity was never resolved in our assays from muscle cells where voltage-activated currents were apparent (Figure 2). Similarly, recordings from tegument-derived vesicles were also negative for responses to PZQ, even though other channels can be resolved (Robertson et al., 1997). But again, it is possible that these membrane vesicles do not capture the diversity of proteins residing within the tegument. The conclusion that PZQ is directly neuroactive is none-the-less consistent with prior attempts that failed to resolve PZQ-evoked currents in recordings from isolated schistosome muscle cells, or the accessible tegument surface. This is especially notable in light of the large, PZQ-evoked conductance of TRPM_PZQ_ which should make this current easy to resolve.

Second, assignment of neuronal identity. Caution is needed in qualifying that TRPM_PZQ_ recordings are made from ‘putative’ neuronal tissue. All labeling was performed using a vital dye rather than a genetically-encoded marker that could unambiguously mark neuronal tissue as used, for example, in genetically tractable models. While *in situ* recordings have long been established with *C. elegans* (Goodman et al., 1998, Qian et al., 2008), progress has been facilitated in this free-living nematode model by various advantages, most notably a well-optimized transgenic toolkit for cell labelling coupled with facile genetic knockout methods (Francis et al., 2003, Goodman et al., 2012). This remains a limitation of the (parasitic) flatworm model as a well-established, routine transgenic methodology is yet to gain traction (but see (Ittiprasert et al., 2023, Weill et al., 2023)). This therefore necessitated the cruder approach of targeting an area of fluorescence signal in a live worm with a recording electrode. Consequently, it is impossible to know with absolute certainty that measurements are truly made from neurons. This is relevant for recordings made from the anterior ganglion area, where no obvious electrical activity was apparent prior to PZQ addition. Even for recordings from the longitudinal nerve cord tissue, where an endogenous oscillatory current was observed (Figure 4), this still cannot unambiguously be attributed to motor neurons, the nerve cord, or the associated nerve plexus. The ability to record this endogenous oscillatory activity does provide opportunity to dissect the underlying ion channels mediating this waveform in future work.

Recognition of both these methodological caveats does not detract from the key advance reported in this study - the definition of a native current evoked by PZQ in a live schistosome, captured at single channel resolution. The properties of this native current are consistent with those of *Sm*.TRPM_PZQ_ measured in heterologous expression systems.

## Acknowledgements

**Author’s Contributions.** EC performed electrophysiological studies in schistosome worms. CMR performed other analyses, including worm isolation, drug treatments and microscopy. JSM wrote the initial draft of the manuscript and supervised this project. All authors worked on revisions and approved the final version of the manuscript.

**Financial Support.** This work was supported by the National Institutes of Health Grant R01-AI145871 (J.S.M.).

**Competing Interests.** The authors declare there are no conflicts of interest.

**Ethical Standards.** Isolation of schistosome worms followed ethical regulations approved by the MCW IACUC committee.

